## Supplementary Material for "Molecular basis for redox control by the human cystine/glutamate antiporter System xc^-^"

**Supplementary Figures 1-9**

### Molecular basis for redox control by the human cystine/glutamate antiporter System xc-.

Joanne L. Parker[1, *, #], Justin C. Deme[2,3,4#], Dimitrios Kolokouris[1], Gabriel Kuteyi[1], Philip C. Biggin[1], Susan M. Lea[2,3,4 *], Simon Newstead[1,5*].

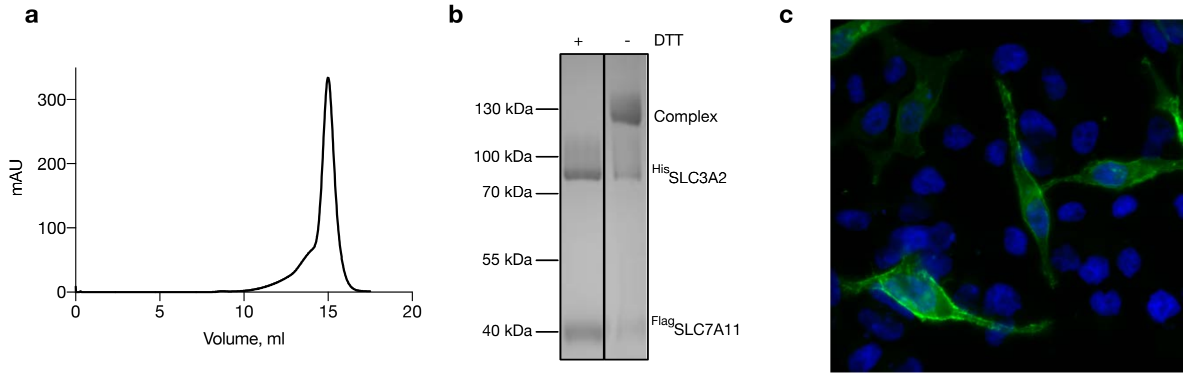

**Supplementary Fig. 1 a**, representative gel filtration trace of SLC7A11/SLC3A2 in LMNG:CHS. **b**, SDS-PAGE analysis of the purified complex in the presence or absence of 1 mM DTT, which dissociates the complex. **c**, Overexpressed GFP tagged SLC7A11 in the presence of SLC3A2 in HeLa cells is targeted mainly to the plasma membrane, Green SLC7A11-GFP, Blue DAPI.

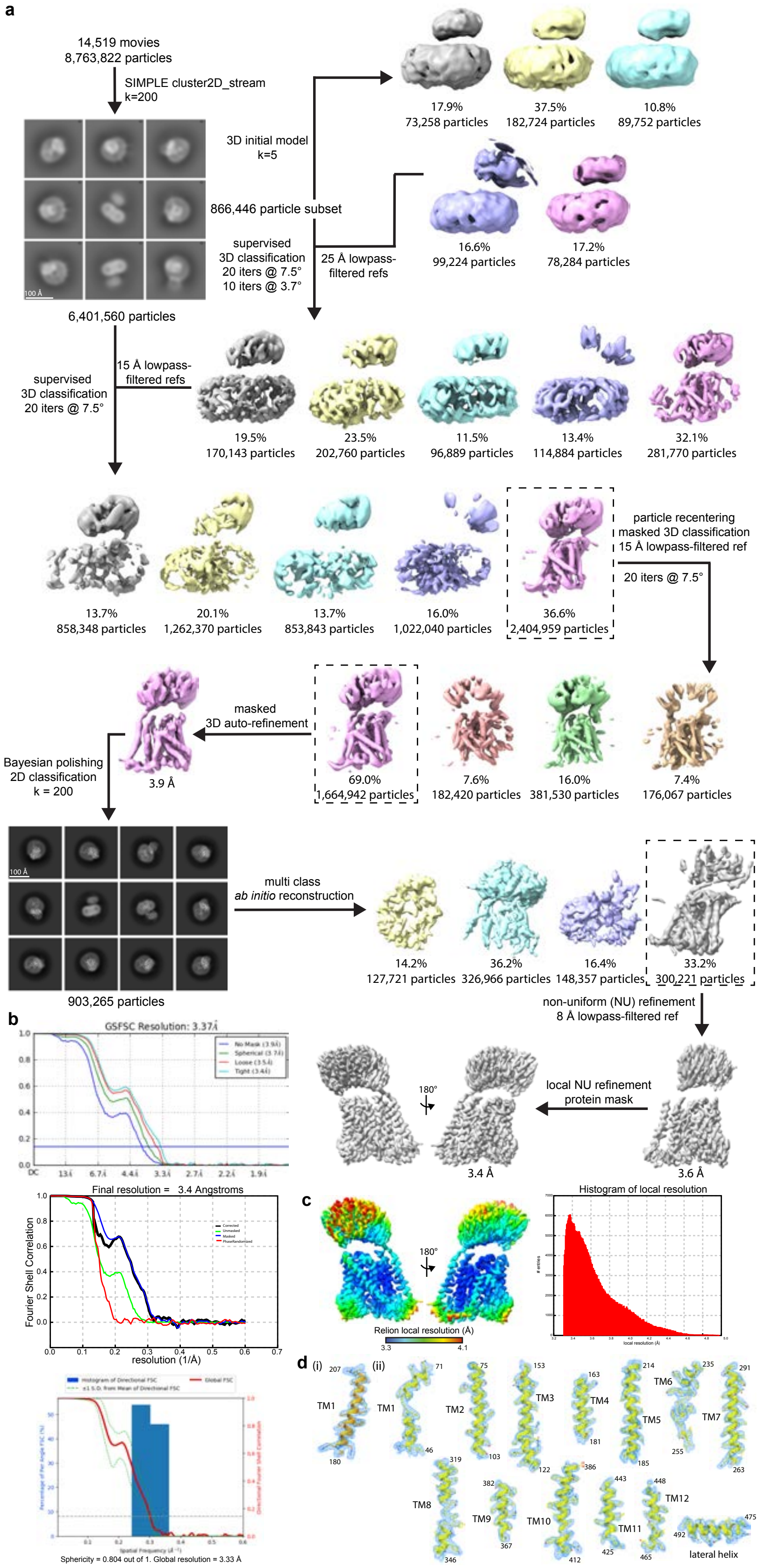

**Supplementary Fig. 2. Cryo-EM processing workflow and local/global map quality for System xc-. a,** Image processing workflow for System $X_C^-$. **b,** Gold-standard Fourier Shell

Correlation (FSC) curves used for global resolution estimates within cryoSPARC (top), RELION (middle), or 3DFSC (bottom). **c,** Local resolution estimation of reconstructed map as determined within RELION. Detergent density omitted for clarity. **d,** Close-up view of map and side-chain density for transmembrane helices and lateral helix. Volume contoured at threshold level of 0.25-0.3.

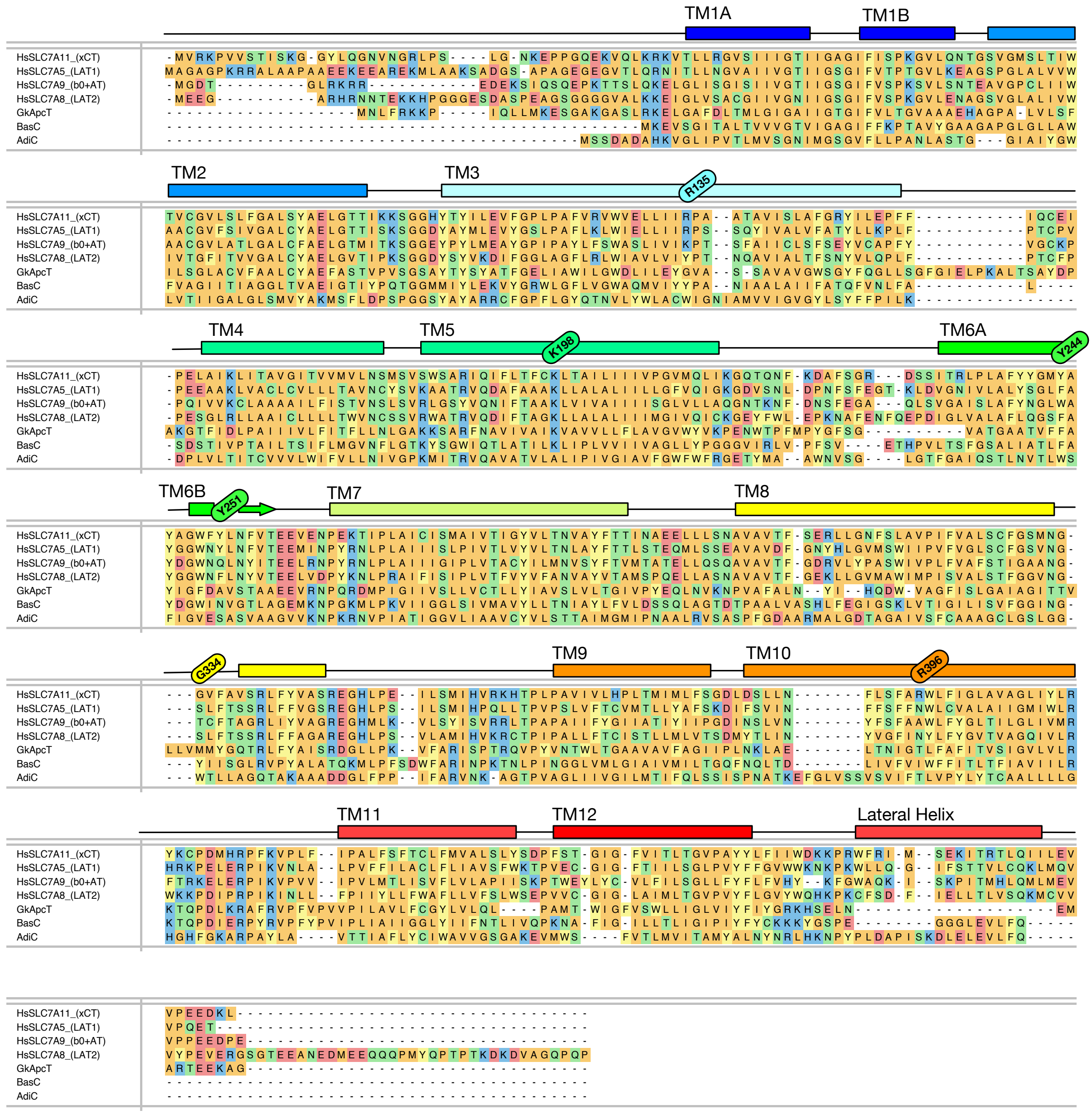

**Supplementary Fig. 3** Sequence alignment of eukaryotic and prokaryotic SLC7 family members with solved structures. Residues with functional importance are highlighted as coloured ovals.

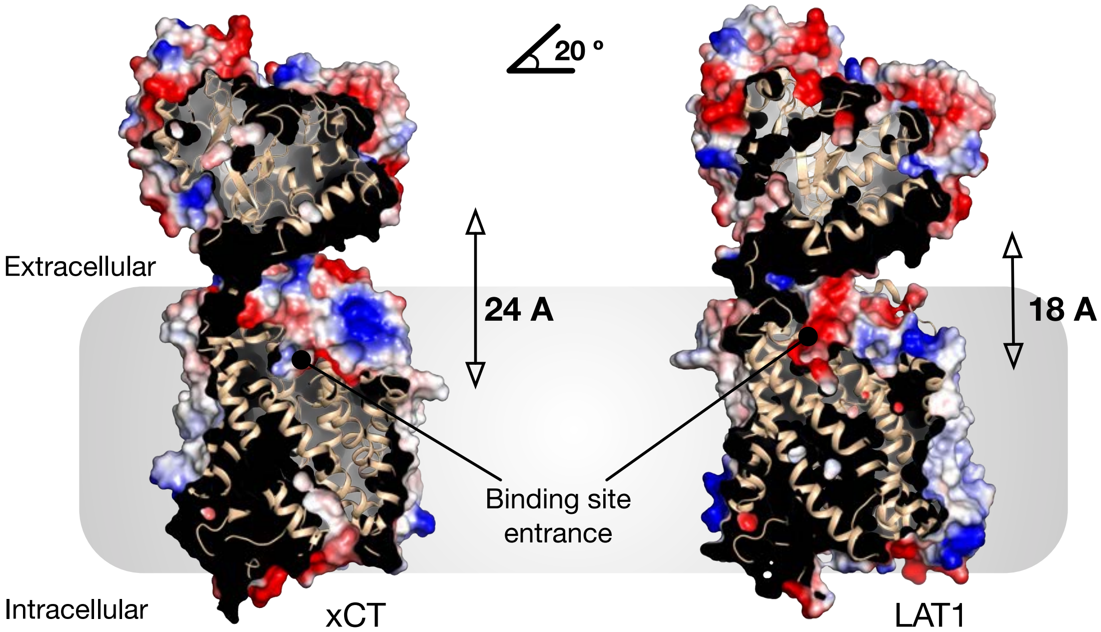

**Supplementary Fig. 4 Structural comparison of the ectodomain of 4F2hc.** The ectodomain is tilted approximately 20 ° away from the transporter domain relative to the position in LAT1, resulting in a wide entrance vestibule to the binding site.

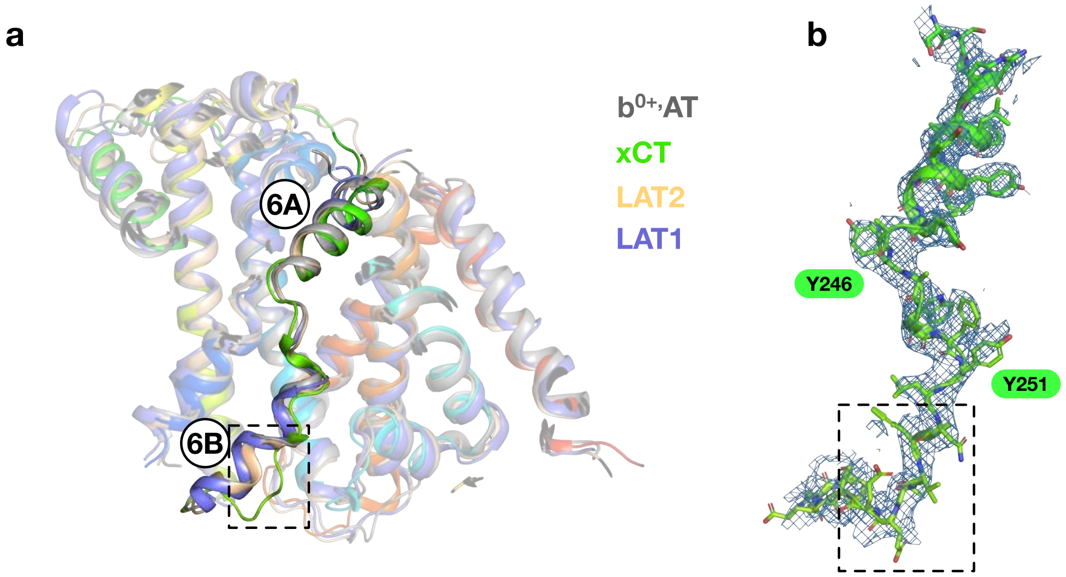

**Supplementary Fig. 5 TM6B in xCT adopts a more disordered structure compared current human SLC7 structures**. **a.** Structural overlay of the human SLC7 transporters highlighting TM6A and TM6B. The structural overlay consists of $b^{0+,}$AT (PDB:6lid), xCT(PDB:7p9v), LAT1 (PDB:6irt) and LAT2 (PDB:6cmi). **b.** Cryo-EM coulombic density for TM6 in xCT is shown (blue mesh), contoured at 8 sigma.

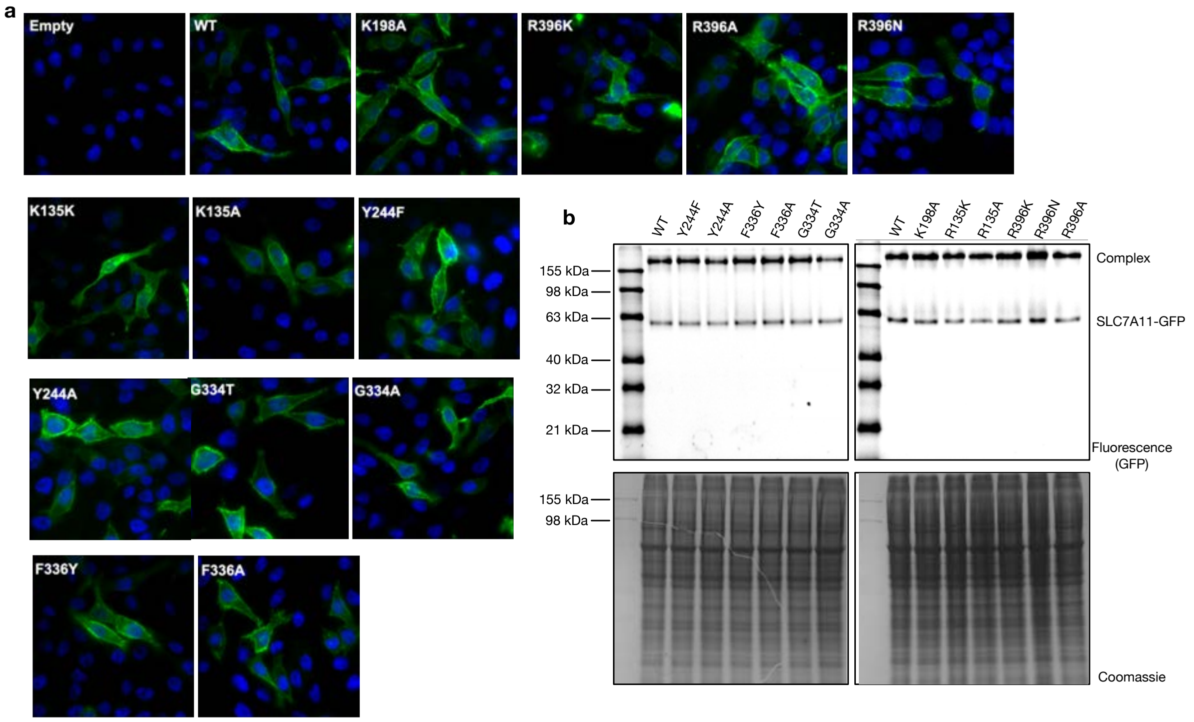

**Supplementary Fig. 6** Expression analysis of the SLC7A11 mutant variants. **a**, Overexpression of the mutant variants of SLC7A11 with a C-terminal GFP tag indicates that all variants are targeted to the plasma membrane, similarly to WT. Green SLC7A11-GFP, Blue DAPI. **b**. In gel fluorescence of C-terminal tagged GFP variants show that all variants are produced in cells to a comparable level (upper panel Fluorescence) (lower panel Coomassie stained gels).

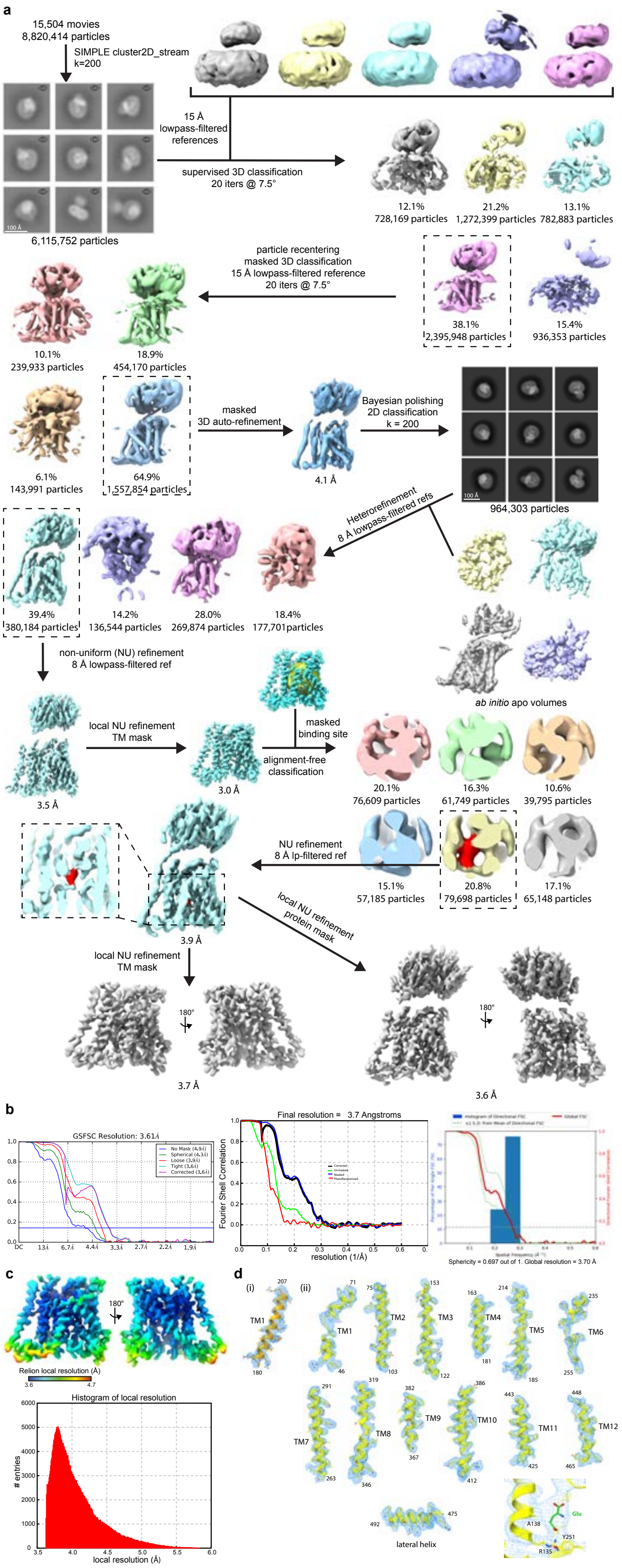

**Supplementary Fig. 7. Cryo-EM processing workflow and local/global map quality for System xc- in complex with glutamate. a,** Image processing workflow for System xc- in complex with glutamate. **b,** Gold-standard Fourier Shell Correlation (FSC) curves used for global resolution estimates within cryoSPARC (left), RELION (middle), or 3DFSC (right). **c,** Local resolution estimation of focused refinement as determined within RELION. Detergent density omitted for clarity. **d,** Close-up view of map and side-chain density for transmembrane helices, lateral helix, and bound glutamate. Helices were contoured at threshold level of 0.17-0.3 and glutamate at threshold level of 0.34.

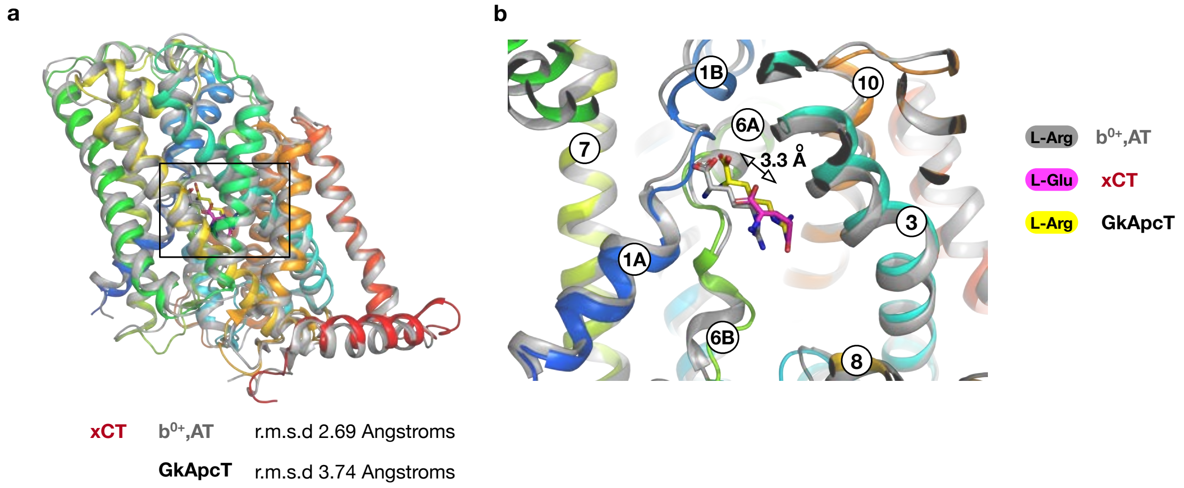

**Supplementary Fig. 8 Comparison of the L-glutamate binding pose in xCT**. **a.** Structural overlay of the xCT-glutamate complex with $b^{0+,}$AT in complex with L-Arginine (PDB:6lid) and GkApcT in complex with L-Arginine (PDB:6f34). For clarity the GkApcT backbone has been omitted. The r.m.s.d. values are given for these structures overlaid onto xCT (PDB:7P9U). **b.** Zoomed in view of the binding site showing the location of L-Arginine in $b^{0+,}$AT and GkApcT compared to xCT.

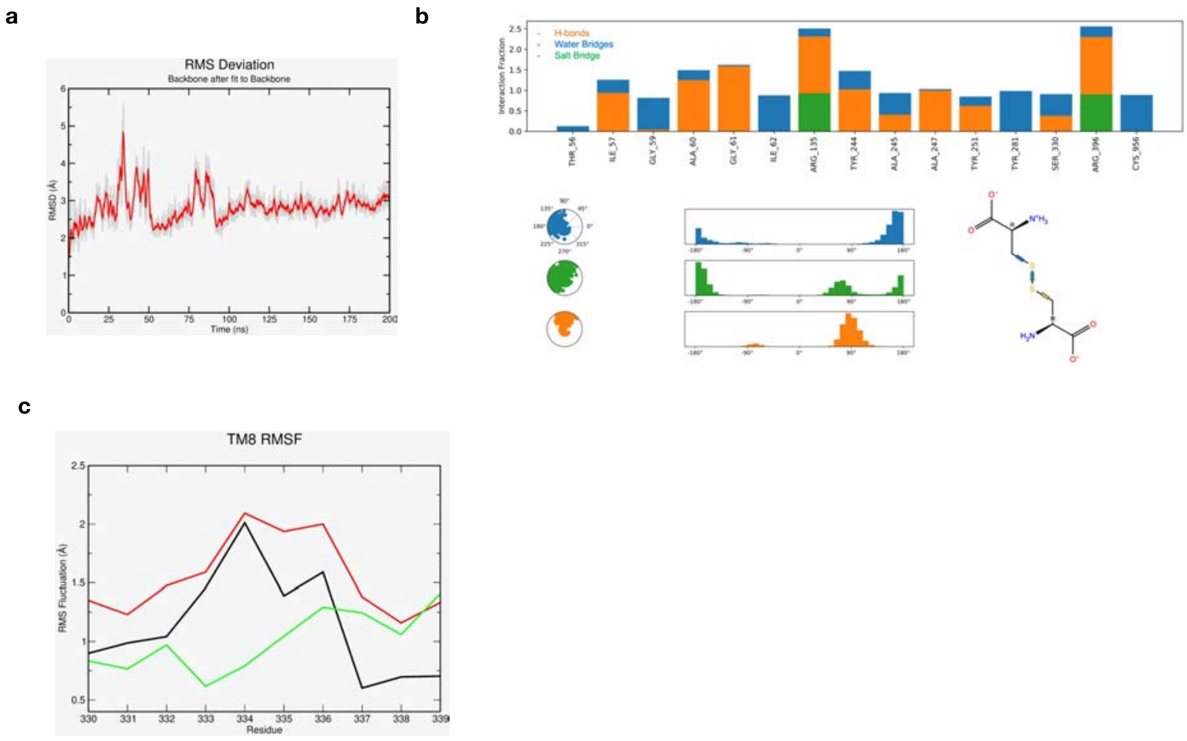

**Supplementary Fig. 9 Molecular Dynamics (MD) simulation analysis of xCT with L-cystine**. **a,** Root-mean-square deviation (in Å) of apo- xCT backbone atoms upon a least squares fit from a 200 ns simulation of the xCT-4F2hc heterodimer. The structure reaches an equilibrium plateau meaning the structure is stable and fit for MD simulation. **b**. Interaction fingerprint of L-cystine bound to xCT binding site residues, calculated as a fraction of the total simulation time from 3 unbiased simulations (3 x 200 = 600 ns). Cystine interactions per residue can take values > 1 (uncapped), thus accounting for more than 1 atoms per residue interacting with cystine in a single frame (orange: H-bonds, blue: water bridge, green: salt bridge interactions. Below, analysis of the cystine dihedrals colour-coded on the cystine chemical structure. Radial plots illustrate the conformational changes of a dihedral over time, and each distribution plot includes a histogram analysis of the dihedrals sampled (bin-width = 10°). **c**, Root-mean-square fluctuation monitoring (in Å) of TM8 residues 330-339 (black: apo- xCT, red: L-Glutamate bound xCT, green: L-cystine bound xCT. When L-cystine is bound, TM8 residues are less mobile and more structured indicating increased helicity in the cystine-bound form.

**Table 1. Cryo-EM data collection, refinement and validation statistics**

| | System $X_C^-$ (EMD-13267) (PDB 7P9V) | System $X_C^-$ + glutamate (EMD-13266) (PDB 7P9U) |
|---|---|---|
| **Data collection and processing** | | |
| Magnification | 105,000 | 105,000 |
| Voltage (kV) | 300 | 300 |
| Electron exposure (e–/Å$^2$) | 59.1 | 59.1 |
| Defocus range (μm) | 0.8 - 2.5 | 0.8 - 2.5 |
| Pixel size (Å) | 0.832 | 0.832 |
| Symmetry imposed | C1 | C1 |
| Initial particle images (no.) | 8,763,822 | 8,820,414 |
| Final particle images (no.) | 300,221 | 79,698 |
| Map resolution (Å) | 3.4 | 3.7 |
| FSC threshold | 0.143 | 0.143 |
| Map resolution range (Å) | 3.3-4.1 | 3.6-4.7 |
| **Refinement** | | |
| Initial model used (PDB code) | None | None |
| Model resolution (Å) | 3.4 | 3.7 |
| FSC threshold | 0.143 | 0.143 |
| Model resolution range (Å) | 3.3-4.1 | 3.6-4.7 |
| Map sharpening *B* factor (Å$^2$) | -81.5 | -74.4 |
| Model composition | | |
| Non-hydrogen atoms | 7286 | 3933 |
| Protein residues | 919 | 502 |
| Ligands | NAG: 8 | GLU: 1 |
| *B* factors (Å$^2$) | | |
| Protein | 67.17 | 90.04 |
| Ligand | 99.12 | 75.62 |
| R.m.s. deviations | | |
| Bond lengths (Å) | 0.005 | 0.003 |
| Bond angles (°) | 0.803 | 0.678 |
| Validation | | |
| MolProbity score | 2.28 | 2.02 |
| Clashscore | 15.39 | 11.18 |
| Poor rotamers (%) | 0.00 | 0.00 |
| Ramachandran plot | | |
| Favored (%) | 88.52 | 92.77 |
| Allowed (%) | 11.48 | 6.83 |
| Disallowed (%) | 0.00 | 0.40 |
